# Potential PGPR properties of cellulolytic, nitrogen-fixing, and phosphate-solubilizing bacteria of a rehabilitated tropical forest soil

**DOI:** 10.1101/351916

**Authors:** Amelia Tang, Ahmed Osumanu Haruna, Nik Muhamad Ab. Majid

## Abstract

In the midst of major soil degradation and erosion faced by tropical ecosystems, rehabilitated forests are established to avoid further deterioration of forest land. In this context, cellulolytic, nitrogen-fixing (N-fixing), and phosphate-solubilizing bacteria are very important functional groups in regulating the elemental cycle and plant nutrition, hence replenishing the nutrient content in forest soil. As other potential plant growth-promoting (PGP) rhizobacteria, these functional bacteria could have cross-functional abilities or beneficial traits that are essential for plants and improve their growths. This study was conducted to isolate, identify, and characterize selected PGP properties of these 3 functional groups of bacteria from tropical rehabilitated forest soils at Universiti Putra Malaysia Bintulu Sarawak Campus, Malaysia. Isolated cellulolytic, N-fixing and phosphate-solubilizing bacteria were characterized for respective functional activities, biochemical properties, molecularly identified, and assessed for PGP assays based on seed germination and indole-3-acetic acid (IAA) production. Out of 15 identified bacterial isolates exhibiting beneficial phenotypic traits, a third belong to genus *Burkholderia* and a fifth to *Stenotrophomonas* sp. with both genera consisting of members from two different functional groups. Among the tested bacterial strains, isolate *Serratia nematodiphila* C46d, *Burkholderia nodosa* NB1, and *Burkholderia cepacia* PC8 showed outstanding cellulase, N-fixing, and phosphate-solubilizing activities, respectively. The results of the experiments confirmed the multiple PGP traits of selected bacterial isolates based on respective high functional activities, root, shoot lengths, and seedling vigour improvements when bacterized on mung bean seeds, as well as presented some significant IAA productions. The results of this study indicated that these functional bacterial strains could potentially be included in future biotechnological screenings to produce beneficial synergistic effects *via* their versatile properties on improving soil fertility and possible crop growth stimulation.

## Introduction

Albeit vast in carbon and biodiversity, tropical ecosystems are facing major deforestations, leading to widespread soil erosion which garnered much recent attention [1–3]. As soil erosion aggravates, the soil loss overrides soil establishment, which results in diminishing soil resources and the ecosystem supports they render [4–5], in particularly manifested as low productivity and farming sustainability on the degraded soils [6]. Planted forest or rehabilitation efforts are undertaken to avoid further deterioration of forest land.

Bacteria has large biodiversity composition in soils, and thus, is major participant of soil principal processes that regulate whole terrestrial ecosystems operations, by means of completing nutrient, and geochemical cycles, including C,N,S, and P [7–9]. In plant litter mass, 20-30% portion are constituted as cellulose, making the latter as most abundant biopolymer [10]. Decomposition of cellulose has been established to be primarily accountable by soil microorganisms, which assimilate the simple sugars resulting from the reduction process of the complex polysaccharides [11–12]. Despite the general ruling capabilities of fungi over bacteria in cellulose decomposition in soils [13–14], these capabilities are also reserved to some, albeit phylogenetically different taxa of bacteria [11]. Previous reports have emphasized bacterial taxa which were recently discovered to possess cellulolytic capabilities, in which they were formerly not known to reduce cellulose [15–16]. To this end, the study of cellulolytic bacterial diversity in tropical soils is warranted.

Nitrogen needs by plant are supported by means of organic compound degradation, atmospheric deposition of N, and biological nitrogen-fixation (BNF), whereby BNF supplies 97% of natural N reserve [17] *via Bacteria* and *Archaea* [18–19]. The N allocation in tropical ecosystems contributed by N-fixing microorganisms (diazotrophs), prevailing mostly in free-living condition, could range from 12.2 to 36.1 kg ha^−1^ year^−1^ [20]. Considering the importance of free-living diazotrophic communities to ecosystem processes, there is a need to broaden our knowledge on their diversity in rehabilitated tropical forests. This is essential because there is dearth of information of this kind.

In most tropical soils which are usually very acidic, P is one of the most commonly deficient macronutrients, owing to the prevalence of typically high Al and/or Fe level, in which P is fixed as insoluble mineral complexes, causing unavailability of P for root uptake [21]. Apart from chemical treatments, microorganisms have also been applied over the years to improve nutrient availability, especially P in soils, as well as alleviating Al toxicity [22]. Furthermore, tropical acid soils support only limited acid-tolerant plant species and microbes. In soils, phosphate-solubilizers are predominated by bacteria over fungi at 1-50% and 0.1-0.5%, of total population, respectively [23]. Bacterial strains consisting of *Pseudomonas, Bacillus*, and *Rhizobium* have been cited to be the dominant phosphate-solubilizers [24]. In addition, *Burkholderia* strains and species which occur in broadly different natural niches have been frequently reported as plant-associated bacteria, albeit their prevalence as free-living in the rhizosphere, as well as epiphytic or endophytic, obligate endosymbionts, or phytopathogens [25]. The phosphate-solubilizing bacteria could release substantial amount of organic acids crucial for fixing Al *via* a chelation process, thus reducing Al toxicity to plant roots in the highly weathered soils [26].

As other potential PGPR, these functional bacteria could have cross-functional abilities or beneficial traits, apart from their primary functional ability such as the production of PGP phytohormones, polysaccharides, and organic acids that are essential for plant growth and development [22]. In addition, IAA which is known to stimulate shoot elongation and/or in particularly root structures by those IAA produced by the bacteria [27], could result in more efficient elemental nutrient acquisition by host plants, leading to higher plant growth and development. To be specific, adoption of functional bacterial strains with multiple functional and PGP traits in crop production would not only promote sustainably increased crop productivity, the practice also offers an option for minimizing costs related to chemical fertilizers application, while at the same time reducing the pollution risks from rampant application of chemical fertilizers.

However, the isolation and characterization of these particular three types of potential functional bacteria from tropical soils of rehabilitated forest remain scarce. Thus, the aims of this study were to: 1) isolate and identify cellulolytic, N-fixing, and phosphate-solubilizing bacteria from a rehabilitated tropical forest soil at Universiti Putra Malaysia Bintulu Sarawak Campus, Malaysia, and 2) characterize cellulolytic, N-fixing, and phosphate-solubilizing bacteria for their respective functional activities, such as IAA production, early plant growth promotion, some phenotypic, and biochemical properties.

## Materials and Methods

### Soil sampling

Soil samples were collected from a rehabilitated forest site at Universiti Putra Malaysia Bintulu Sarawak Campus through random sampling design using aseptic techniques near rhizospheric zones of planted trees. Soil samples were collected at 0-10 cm depth using an auger. Prior to soil sampling, any litter at the surface was manually removed [28]. The plastic bags with soil samples were transported to the laboratory at Universiti Putra Malaysia Bintulu Sarawak Campus for analysis. The soil samples were homogenized or mixed thoroughly to make composite (consolidated) samples, and surface organic materials, litter, roots, rocks, and macro fauna were removed. Soil samples were kept in tightly sealed plastic bags and stored at 4 °C (for less than 3 days before microbial analysis) to keep the soil moist, and to preserve biological properties.

### Isolation of free-living nitrogen-fixing bacteria

Ten-fold serial dilution technique was used to determine the CFU number of the culturable N cycle functional bacteria. Bacterial colonies were enumerated on N-free malate (Nfb) medium [29] with some modifications containing per litre: 0.4 g of KH_2_PO_4_, 0.1 g of K_2_HPO_4_, 0.2 g of MgSO_4_.7H_2_O, 0.1 g of NaCl, 0.02 g of CaCl_2_, 0.01 g of FeCl_3_, 0.002 g of MoO_4_Na.2H_2_O, 5.0 g of sodium malate, 5 mL of bromothymol blue with 0.5% alcohol solution and 15.0 mL of agar. The whole media content were adjusted to pH 7 before being sterilized in autoclave (121 °C for 20 minutes). Approximately 20 mL of media was placed in each petri dish before being incubated at 30 °C after triplicate inoculation. The colonies that fix N_2_ alkalinized the culture medium (blue) due to ammonia production. Some colonies exhibiting large and instant blue halo on the medium were isolated, purified on new Nfb media, sub cultured on agar slants, and stored at 4 °C for further studies.

### Isolation of phosphate-solubilizing bacteria

Cells and spores were detached from soil surface by shaking at 150 rpm a mixture of 5 g of fresh soil and 100 mL of sterile distilled water for 24 hr. Supernatant was ten-fold serially diluted in sterile distilled water and 0.1 mL aliquots in triplicate were spread over Petri dishes containing differential growth media for phosphate-solubilizing bacteria. This isolation technique is based on the formation of halos around the colonies of bacteria capable to solubilize a calcium phosphate insoluble compound [30–31]. The inoculated plates were incubated at 30 °C. Culture media employed to differentially isolate phosphate-solubilizing bacterial strains consisted of variations of a chemically defined medium [32–33] with some modifications. All media contained per litre: 50 mL of a salts solution (MgCl_2_·6H_2_O, 100 g L^−1^; MgSO_4_·7H_2_O, 5 g L^−1^; KCl, 4 g L^−1^; (NH_4_)_2_SO_4_, 2 g L^−1^), 10 g of sucrose as C source, and 2.5 g of tricalcium phosphate, Ca_3_(PO_4_)_2_, as an insoluble source of phosphate. Six mL L^−1^ of a pH indicator dye (bromophenol blue, 0.4% ethanol solution) was added to the media [30]. For solidification, 20 g L^−1^ of agar was added and pH of medium was adjusted to 7. Phosphate-solubilizing bacterial colonies exhibiting large and instant halos on the media were isolated, further purified on new media, sub cultured on agar slants, and stored at 4 °C for further studies.

### Isolation of cellulolytic bacteria

This procedure was modified from those as described by Hendricks et al. [34]. The cellulolytic bacteria was enumerated by triplicate spread plate inoculation onto cellulose-Congo red agar with 10-fold sterile tap water dilutions of soil, and incubated at 30 °C. The cellulose-Congo red agar consisted of 0.50 g of K_2_HPO_4_, 0.25 g of MgSO_4_, 1.88 g of cellulose microgranular powder, 0.20 g of Congo red, 5.00 g of agar, 2.00 g of gelatin, 100 mL of soil extract, and 900 mL of tap water (used to provide essential trace elements for soil bacteria), with pH adjusted to 7. Following incubation, the colonies exhibiting zones of clearing were counted. Some colonies exhibiting large and instant zones of clearing were isolated, further purified on the same media, sub cultured on agar slants, and stored at 4 °C for further studies.

## Plate assays for cellulose-hydrolyzing, nitrogen-fixing, and phosphate-solubilizing activities

### Plate assay for cellulase activity by cellulolytic isolates

Based on clear zones formation on cellulose-Congo red agar as modified from Hendricks et al. [34], cellulolytic bacteria were isolated and evaluated for their cellulose-hydrolyzing index values using same medium with following composition: K_2_HPO_4_, 0.50 g L^−1^; MgSO_4_, 0.25 g L^−1^; cellulose powder, 1.88 g L^−1^; Congo red, 0.20 g L^−1^; agar, 15.00 g L^−1^; gelatine, 2.00 g L^−1^; 100 mL of soil extract, and 900 mL of tap water to supply essential trace elements. The medium was maintained at pH 7. Following incubation at 30 °C for over 10 day-period, cellulolytic isolates were assessed for their selective abilities index values based on calculation of ratio of total diameter (colony + halo or clearing zone around colonies) and colony diameter [35].

### Plate assay for nitrogen-fixation activity by nitrogen-fixing isolates

Nitrogen-fixing bacteria were cultured on N-free malate (Nfb) medium modified from Döbereiner and Day [29] and incubated at 30 °C. Bacterial colonies with blue halo formation were isolated and characterized using the same media, after which blue halo index values were calculated. The Nfb medium consisted of: KH_2_PO_4_, 0.4 g L^−1^; K_2_HPO_4_, 0.1 g L^−1^; MgSO_4_.7H_2_O, 0.2 g L^−1^; NaCl, 0.1 g L^−1^; CaCl_2_, 0.02 g L^−1^; FeCl_3_, 0.01 g L^−1^; Mo.O_4_Na.2H_2_O, 0.002 g L^−1^; sodium malate, 5 g L^−1^; agar, 15 g L^−1^; bromothymol blue (0.5% in alcohol solution), 5 mL L^−1^. The medium was adjusted to pH 7. Following incubation at 30 °C for over 10 day-period, all N-fixing isolates were assessed for their selective abilities index values based on calculation of ratio of total diameter (colony + halo or clearing zone around colonies) and colony diameter [35].

### Plate assay for phosphate-solubilization activity by phosphate-solubilizing isolates

Phosphate-solubilizing bacterial isolation technique is based on the formation of clear halos around the colonies capable to solubilize calcium phosphate [30–31]. Media containing insoluble phosphate [32–33] with some modifications were used to isolate and assess the phosphate-solubilizing bacterial activities. Its composition was: 50 mL L^−1^ of a salts solution (MgCl_2_·6H_2_O, 100 g L^−1^; MgSO_4_·7H_2_O, 5 g L^−1^; KCl, 4 g L^−1^; (NH_4_)_2_SO_4_, 2 g L^−1^); agar, 15 g L^−1^; 10 g L^−1^ of sucrose as C source, and 2.5 g L^−1^ of tricalcium phosphate, Ca_3_(PO_4_)_2_, as an insoluble source of phosphate, and content was adjusted to pH 7 before being autoclaved. Six mL L^−1^ of a pH indicator dye (bromophenol blue, 0.4% ethanol solution) was also added to the media [30]. Following incubation at 30 °C for over 10 day-period, phosphate-solubilizing bacterial isolates were assessed for their selective abilities index values based on calculation of ratio of total diameter (colony + halo or clearing zone around colonies) and colony diameter [35].

## Quantitative assays for functional activities

### Quantitative assay for cellulase activity

For quantitative determination of cellulase activity, cellulolytic isolates were previously inoculated in nutrient broth, shaken at 120 rpm at 30 °C for 16 hr and size of the culture as preinoculum for cellulase assay was fixed to OD_600_=0.5 (10^6^ to 10^7^ cfu mL^−1^). One mL of each isolate preinoculum was seeded in 40 mL cellulose broth as basal medium containing cellulose as sole C source. The composition of cellulose broth per litre was: 5.0 g L^−1^ cellulose microgranular, 2.5 g L^−1^ NaNO_3_, 1.0 g L^−1^ KH_2_PO_4_, 0.6 g L^−1^ MgSO_4_·7H_2_O, 0.1 g L^−1^ NaCl, 0.1 g L^−1^ CaCl_2_·6H_2_O, 0.01 g L^−1^ FeCl_3_, 2.0 g L^−1^ gelatin, and 0.1 g L^−1^ yeast extract with pH 7 adjusted using diluted NaOH as modified from Lu et al. [36]. Inoculated broth was incubated at 30 °C under agitation at 180 rpm and CMCase activity was analyzed for up to 5 days. For CMCase activity, 0.5 mL culture filtrate from broth as crude enzyme source, or supernatant cellulase, also specified as CMCase, was incubated with 1 mL of 1% carboxymethylcellulose in 0.1 M citrate buffer, pH 5 at 40 °C in agitation for 30 min. The amount of reducing sugars (glucose) was measured using Somogyi-Nelson method [37–38] at 520 nm. One unit of enzyme activity (IU mL^−1^) is equivalent to the amount of enzyme per mL culture filtrate that liberates 1μg of glucose per minute [39].

### Quantitative assay for *in vitro* biological nitrogen-fixation activity

The procedure for quantitative estimation of BNF activity was modified from those described by Soares et al. [40]. Each N-fixing isolate was grown as a preinoculum in nutrient broth under agitation at 120 rpm at 30 °C for 16 hr. Optical density was used to adjust the inoculum size [OD_600_= 0.5 is equivalent to 10^6^ to 10^7^ colony forming units (cfu) mL^−1^]. Inoculum was transferred (100 μL) to new media, NFb broth (containing similar composition as Nfb medium except without agar). The inoculated media were incubated at 30 °C under constant agitation for 72 hr for N_2_ fixation quantification. A control medium was also included by using 100 μL nutrient broth as inoculant. The amount of N_2_ fixation was measured by sulfur digestion and distillation with NaOH 10 mol L^−1^, as described by Bremner and Keeney [41] and done in triplicates. The samples were digested with 2 mL concentrated H_2_SO_4_ (*d* = 1.84), 1 mL H_2_O_2_ 30% and 0.7 g digestion mixture (100 g Na_2_SO_4_ + 10 g CuSO_4_.5H_2_O + 1 g selenium) at 350-375 °C. The products of digestion were distillated with 10 mL NaOH 10 mol L^−1^, and the ammonia trapped in 10 mL of 2% boric acid was measured by titration with H_2_SO_4_ 0.0025 mol L^−1^.

### Quantitative assay for inorganic phosphate-solubilization activity

Quantitative estimation of inorganic phosphate-solubilization was done with some modifications from Delvasto et al. [42]. Preinoculum cultures were previously prepared by inoculating phosphate-solubilizing isolate in nutrient broth under agitation at 120 rpm at 30 °C for 16 hr. Inoculum size was adjusted to OD_600_=0.5 (10^6^ to 10^7^ cfu mL^−1^). Prior to the assay, preinoculum cultures were washed 3 times and resuspended using saline solution 0.85% NaCl, and one mL isolate suspension was inoculated into 100 mL liquid medium containing insoluble calcium phosphate as in solid medium, except without agar. Inoculated broth was incubated at 30 °C under agitation at 180 rpm for up to 5 days. Saline solution without inoculant in the broth was left as control.

Soluble P content in culture filtrates was determined by the molybdenum blue method [43]. Appropriate amount of aliquots from samples containing 1-50 μg mL^−1^ P were pipetted into 50 mL volumetric flasks containing approximately 15 mL distilled water and 8 mL ascorbic acid antimony sulfomolybdo working reagent (as color developing solution to form blue color complex in the present of P from samples and/or standard solutions), swirled to mix, and filled to the mark with distilled water. A series of standard P solutions (KH_2_PO_4_) aliquots were pipetted into a set of volumetric flasks containing the same amount of color developing solutions to make up the range of expected P concentrations in the culture filtrates, swirled to mix, and made up to the mark with distilled water. A blank solution equal to the aliquot size of sample was also included to each flask of standard solutions. A standard curve was plotted using UV-vis spectrophotometer (Unico) at 882 nm. The absorbance of samples were measured by means of the standard curve using the same wavelength and converted into P concentrations. Values of soluble P are expressed as μg mL^−1^. The final pH of liquid medium was measured using a pH meter. All values of pH and P concentrations in samples were measured in triplicates.

### Morphological and physiological characterization of selected isolates

Selected isolates were each studied for morphological properties, which included color, size, and colony characteristics such as form, margin, and elevation on nutrient agar. The spore staining procedure, done according to Schaeffer-Fulton method and standard Gram staining procedure were conducted as described by Cappuccino and Sherman [44], both with some modification on fixation, whereby bacterial smears were fixed using methanol, without heat fixation. For each procedure, the smears were finally examined under oil immersion at total magnification power of 1000 times. The morphology of each bacterial colony such as the color, shape, and arrangement of the stained cells were recorded. Any presence of endospores for each studied bacterial isolate was also noted.

The biochemical tests consisting of several enzymatic, physiological activities, or differential growth tests were carried out. Citrate utilization was determined on Simmons citrate agar; hydrogen sulfide production, along with lactose and glucose fermentation were determined on Kligler Iron Agar; Gram negative strains and lactose fermentation were detected using Mac Conkey agar; gelatinase production, starch hydrolysis, casein proteolysis, lipase production, and indole production were each determined using nutrient gelatine, starch agar, skim milk agar, Tween 80 agar and BD’s Indole Reagent Droppers (modified Kovacs’ reagent) *via* standard tube method, respectively. Catalase test was done using 3% hydrogen peroxide solution. Motility test was conducted on Biolife’s Motility Medium whereas nitrate reduction test was done using the test kit by Fluka. The aforementioned physiological and biochemical test results were processed and compared with the results of systematic bacteriology identification according to Bergey’s Manual of Determinative Bacteriology and Bergey’s Manual of Systematic Bacteriology [45–46].

## Molecular method for bacterial identification

### Deoxyribonucleic acid extraction

Bacterial genomic DNA was extracted using sodium dodecyl sulfate (SDS) method. Bacterial inoculum from streaked culture plate overnight was recultured overnight in 1.5 mL nutrient broth in capped tubes with one control (broth without inoculum) in the water bath shaker at 30 °C. The 1.5 mL cell suspension was centrifuged at 13040 *g* for 3 min at 4 °C. After removing the supernatant, the cells were resuspended or washed with 180 μL TE buffer (10 mM Tris, 1mM EDTA, pH 8.0) by vortex-mixing and incubated on ice for 5 min. Seventy five μL of 2% SDS solution was added, followed by invert-mixing step to lyse the bacterial cells and incubation on ice for 5-10 min. Then 250 μL of 3 M potassium acetate was added, followed by gentle invert-mixing, and incubation on ice for 5 min. The sample was subsequently centrifuged at 13040 *g* for 5 min at 4 °C, with special attention on the cells pellet formation at the bottom of the tube. The supernatant containing bacterial DNA was carefully transferred to a clean 1.5 mL tube on ice. Absolute ethanol was then added at approximately two times the volume of the supernatant on the same tube, invert-mixed, and followed by centrifugation at 12000 *g* for 10 min at 4 °C.

Later, supernatant was discarded and 1 mL of 75% ethanol was added to the DNA pellet for washing step. DNA pellet was resuspended by pipette-mixing the suspension in repeated gentle motion, followed by centrifugation at 12000 *g* for 10 min at 4 °C. The supernatant was discarded with extra care taken at this stage so as not to disrupt the DNA pellet while at the same time ensuring the entire removal of ethanol residues from the pellet and within the tube itself. The pellet was air-dried for about 10 to 20 min, till no ethanol residue is visible. Finally, 20 μL of TE buffer (10 mM Tris, 1mM EDTA, pH 8.0) was added, pipette-mixed gently, and stored at −20 °C.

### Polymerase chain reaction amplification of extracted DNA

Amplification of DNA extraction products was performed in a final volume of 25 μL each. For each reaction, 1.25 μL 0.5 μM for each of both primers (16S-F and 16S-R1), 2.5 μL 25m*M* MgCl_2_, 2.5 μL 10X reaction buffer [(NH_4_)_2_SO_4_], 0.5 μL working stock dNTPs or dNTP mix combined to give a final concentration of 0.2 m*M* each, 0.25 μL of *Taq* DNA Polymerase (5 U/μL), 15.75 μL of sterilized distilled water and 1 μL of template DNA. Primers 16S-F and 16S-R1 had the sequences of 5’- AAGAGTTTGATCATGGCTCA-3’ and 5’- TAAGGAGGTGATCCAACCGCAGGTTC-3’, respectively. The PCR was performed in a thermal cycler (Bioer XP Cycler) using cycling conditions consisting of an initial denaturation at 95 °C for 3 min, followed by 35 cycles consisting of denaturation at 94 °C for 30 s, annealing at 50 °C for 30 s, and extension at 72 °C for 1 min. A final extension was performed at 72 °C for 5 min. A blank that contained all the components of the reaction mixture without the DNA template was included as a control. The PCR products were analyzed by running on 1% agarose gel electrophoresis.

### Agarose gel electrophoresis

A 1% (w/v) agarose gel was prepared by mixing 0.25 g agarose and 25 mL of 1X TAE buffer. The solution was heated gently in a microwave oven until the agarose was fully dissolved. The gel was allowed to cool to the touch before being poured to the gel cask with the well comb in place. The gel was allowed to solidify for 20-30 minutes: before removing the comb and mould barriers, placed onto electrophoresis tank, and 1X TAE buffer solution was poured evenly into the electrophoresis tank untill it just submerged the surface of the gel. The samples were each prepared by combining 5 μL of PCR products with 3 μL of 6x loading dye and 1 μL of EZ Vision dye before being loaded into the performed wells. One μL of DNA marker or ladder bearing 1.5 kb size was mixed well with 2 μL of 6x loading dye, 1 μL of EZ Vision dye, and 5 μL of 1x TAE buffer solution before being loaded into assigned well. The gel was then run at 90 V for about an hour. The EZ Vision dye-stained fragments were visualized and photographed with UV light in a gel imager (AlphaInnotech Fluorchem). Any PCR bands obtained for the specific isolates were recorded and the gel image was captured. Once PCR bands were sighted on the gel image, preparations of the samples were done which included PCR scale-up to 50 μL per reaction, and purification of PCR products prior to sending them out for outsourced sequencing service.

### Purification of PCR products for sequencing

Direct precipitation is a general method used for purification of PCR products Fifty μL of PCR product was transferred to a 1.5 mL tube. Sodium acetate (pH 8) at 0.1 volume or equivalent to 4 μL was added into the tube, followed by absolute ethanol (99%) at 2.5 volume or equivalent to 100 μL. The tube was inverted a few times for homogeneous mixing and kept in ice for 30 min. The tube was centrifuged at 13,040 *g* for 15 min at 4 °C. Supernatant was decanted and 200 μL of 75% ethanol was added into the tube, followed by vortex-mixing for 1 min. The tube was centrifuged at 13,040 *g* for 5 min at 4 °C. The pellet was air-dried until no traces of ethanol could be seen. Fifteen μL of sterile distilled water was added to resuspend the pellet by tapping. The concentrated PCR product was short-spun and ready to be sent for sequencing. The quantification or estimation of the concentration of the concentrated PCR products was done by using the gel imager software (FluorChem 5500) which was within the range of 5 to 100 ng per μL.

### Sequencing of 16S rDNA

Determination of the 16S rDNA nucleotide sequences was done by transferring the purified PCR products containing the concentrated amplified DNA, along with primer R2 to First BASE Laboratories Sdn Bhd, a life sciences-based research company located in Peninsular Malaysia.

### Identification of bacterial sequences

The sequences obtained for selected bacterial isolates were scanned manually for any errors on the bases interpretation provided by the base sequencing service which consisted primarily of adenine (A), guanine (G), cytosine (C), and thymine (T) bases. The sequence bases were analyzed using Sequence Scanner Software v1.0 which was registered under Applied Biosystems^®^. Each of the corrected sequences was searched on BLAST website which compared nucleotide or protein sequences to sequence databases and calculated statistical significance of matches. The BLAST website was available at http://blast.ncbi.nlm.nih.gov/Blast.cgi. The closest matches of top-scoring sequences from the BLAST databases were used to presume the identity or closest identities of the particular bacteria. In the case of multiple identities produced based on sequence matches alone, biochemical properties offered phenotypic information necessary for further comparison in order to finalize each isolate to only one most presumed identity.

## Plant growth-promoting assays

### Germination assays of maize and green gram seeds

Phytotoxic assay for 15 selected bacterial inoculants on green gram seeds were done *via* germination check of both root and shoot of the germinating seeds prior to pot experiment. Seeds of green gram (*Vigna radiata*) were obtained from a local market. Prior to the assay, the seeds were sorted for uniformity of size, and all visibly damaged seeds were discarded. The remaining seeds were washed with tap water and surface sterilized in 0.1% sodium hypochlorite solution for 5 min before rinsing at least three times with sterile distilled water to thoroughly remove sterilants. The seeds were then soaked in broth containing 16 hr cultured bacterial isolates according to treatments for 2 hr at 30 °C. The seed treatments consisted of 15 selected bacterial isolates. Two control treatments were included whereby first control was designated as seeds treated with sterile distilled water, and another control treatment with sterile nutrient broth.

The treated green gram seeds were placed on petri dishes containing sterilized moist sheets of Whatman filter paper per treatment. Each treatment was based on 100 seeds of each plant type. The seeds were incubated for up to 7 days at about 30 °C and moisture content of growth support was maintained with an equal volume of autoclaved distilled water on daily basis. The number of germinated green gram seeds was firstly counted on third and fourth day of incubation, respectively, and followed by second counting and measurement of shoots and roots or radicle length for both plants on seventh day. Percentage of germinated seeds and vigor index were calculated. Vigor index was calculated as the sum of mean shoot and root length multiplied by percentage of germination [47]. All experimental sets and apparatus used were sterilized where necessary.

### Indole-3-acetic acid production assay

Some selected bacterial isolates were subjected to *in vitro* colorimetric determination of IAA production using Salkowski’s reagent (2% of 0.5 M FeCl_3_ in 35% perchloric acid) as modified from Asghar et al. [48]. Based on seed germination test results, 7 bacterial isolates were selected for IAA production assay. Each selected isolate was grown to exponential growth phase (OD ≥ 1.0) within 16 hr to be used as pre-inoculum for the bioassay. The inoculum was adjusted to OD_600_=0.5 (10^6^ to 10^7^ cfu mL^−1^) before inoculation at 500 μL into 50 mL of nutrient broth without addition of tryptophan and incubated at 30 °C under agitation at 100 rpm for up to 48 hr. Non inoculated broth was included as control. Each culture was centrifuged at 10,000 rpm for 10 min at 4 °C.

One millilitre of culture supernatant was mixed with 2 mL of Salkowski’s reagent in a test tube and kept in the dark for 20-25 min before measuring absorbance using a UV-Vis spectrophotometer at 535 nm. The assay for each isolate was conducted in triplicates. Prior to samples measurement, a series of IAA standards prepared within the range of 0 to 100 μg mL^−1^ reacted with Salkowski’s reagent were measured to construct a standard linear graph as a reference to the samples’ IAA concentration values. Any appearance of reddish to pinkish color in the solution indicated IAA presence in the cultured medium and depending on the color intensity of the test solutions, coloring responses were classified into two; pink type (pale pink to deep pink), and red type (dark red to red).

### Data analysis of bioassays

Analysis of Variance (ANOVA) was used to test for significance differences of measurements of each bioassay whereas Tukey’s Test was employed for the significance difference among different treatments at p ≥ 0.05. The Statistical Analysis System (SAS Ver. 9.2) was used for the analysis.

## Results and Discussion

### Functional activity assays of respective isolates

In this study, 14 out of 62 cellulolytic, 12 out of 39 N-fixing, and 8 out of 36 phosphate-solubilizing bacterial isolates obtained from rhizospheres of both planted and indigenous tree species at the rehabilitated forest were screened based on their functional activities on their respective selective media and distinct morphology for further characterization. Among these 3 functional groups, phosphate-solubilizing group had the least number of characterized isolates due to the fact that phosphate-solubilizing bacterial isolates tend to lose their ability after successive transfers or subcultures on agar medium as reported previously [49]. Diazotrophic isolates also encountered similar phenomenon whereby upon successive cultivation, some isolates failed to develop blue halo or unable to grow on N-free media. da Silva et al. [50] has reported that a diazotrophic isolate lost its characteristic pellicle growth upon successive cultures which made its nitrogenase activity assessment not possible. However, none of the cellulolytic isolates lost their functional ability.

Cellulolytic microorganisms play major role in the synthesis of cellulases crucial for the breakdown of complex polysaccharides into simple sugars. There was no specific correlation between cellulose-hydrolyzing index and cellulase (CMCase) activity for the 14 tested cellulolytic strains (Table 1).

**Table 1.** Cellulose-hydrolyzing index and cellulase (CMCase) activities (U mL^−1^ cell-free culture broth after 5 days at 30 °C, 180 rpm, and medium pH 7) of cellulolytic isolates.

| Isolates | Activity Index | Cellulase Activity (U mL<sup>-1</sup>) |
| --- | --- | --- |
| | Mean value ± S.E. | |
| **C14** | 5.25<sup>bc</sup> ± 1.50 | 0.016<sup>f</sup> ± 0.0015 |
| **C39d** | 5.10<sup>bc</sup> ± 0.44 | 0.015<sup>f</sup> ± 0.0017 |
| **C41d** | 5.29<sup>bc</sup> ± 0.88 | 0.078<sup>b</sup> ± 0.0012 |
| **C42d** | 8.52<sup>ab</sup> ± 1.93 | 0.077<sup>b</sup> ± 0.0007 |
| **C46d** | 10.41<sup>a</sup> ± 0.43 | 0.089<sup>a</sup> ± 0.0009 |
| **C22b** | 10.31<sup>a</sup> ± 0.62 | 0.035<sup>e</sup> ± 0.0006 |
| **C34c** | 4.57<sup>c</sup> ± 3.24 | 0.047<sup>d</sup> ± 0.0010 |
| **C45d** | 5.93<sup>bc</sup> ± 1.23 | 0.059<sup>c</sup> ± 0.0012 |
| **C4** | 3.59<sup>c</sup> ± 0.14 | 0.079<sup>b</sup> ± 0.0015 |
| **C25b** | 3.70<sup>c</sup> ± 0.79 | 0.078<sup>b</sup> ± 0.0012 |
| **C29b** | 5.57<sup>bc</sup> ± 0.70 | 0.080<sup>b</sup> ± 0.0007 |
| **C40d** | 4.63<sup>c</sup> ± 0.61 | 0.077<sup>b</sup> ± 0.0007 |
| **C43d** | 4.60<sup>c</sup> ± 2.26 | 0.078<sup>b</sup> ± 0.0009 |
| **CB15** | 5.54<sup>bc</sup> ± 0.36 | 0.060<sup>c</sup> ± 0.0010 |

S.E. is standard error of the mean.
Means with same letter are not significantly different at $p \geq 0.05$ within columns using Tukey's test.

Among the 14 cellulolytic isolates assessed, isolates C46d and C42d consistently showed higher cellulolytic activity based on both cellulose-hydrolyzing index (10.41 and 8.52, respectively), and cellulase activity (0.089 U mL^−1^ and 0.077 U mL^−1^, respectively). The lowest cellulose-hydrolyzing index and cellulose activity were in the range of 1.33-3.73 U mL^−1^ and 0.015-0.016 U mL^−1^, respectively. The cellulase (CMCase) activities of the cellulolytic isolates in this present study were comparable to those of bacterial strains isolated from soil at different culture conditions as reported by Sethi et al. [51]. The cellulase (CMCase) activity demonstrated by C46d was especially higher than those reported for *Bacillus amyloliquefaciens* SS35 [52] and *Bacillus pumilus* EB3 [53] at 0.079 U mL^−1^.

For nitrogenase activity assessment, there was positive relationship (r=0.63, p ≤ 0.05) between N-fixing index values and BNF amount. Among the 12 diazotrophic isolates evaluated, NB1 exhibited the most active N-fixing activity as demonstrated by the consistent highest means of both activity index and nitrogenase activity quantification at 10.82 and 2.21 μg N mL^−1^ day^−1^, respectively (Table 2).

**Table 2.** Nitrogen-fixation index and biological nitrogen-fixation quantification of diazotrophic isolates.

| Isolate | Activity Index | Quantitative activity ($\mu\text{g N mL}^{-1} \text{ day}^{-1}$) |
| --- | --- | --- |
| | Mean value ± S.E. | |
| NB1 | 10.82<sup>a</sup> ± 0.26 | 2.21<sup>a</sup> ± 0.34 |
| NB4 | 1.84<sup>e</sup> ± 0.05 | 1.11<sup>cd</sup> ± 0.27 |
| NB5 | 2.03<sup>e</sup> ± 0.06 | 0.65<sup>cd</sup> ± 0.00 |
| NB7 | 1.97<sup>e</sup> ± 0.05 | 1.23<sup>bc</sup> ± 0.24 |
| NB8 | 1.80<sup>e</sup> ± 0.05 | 0.47<sup>d</sup> ± 0.00 |
| NC9 | 4.63<sup>c</sup> ± 0.12 | 0.84<sup>cd</sup> ± 0.00 |
| NB10 | 2.35<sup>e</sup> ± 0.32 | 1.01<sup>cd</sup> ± 0.19 |
| NC11 | 1.95<sup>e</sup> ± 0.04 | 0.44<sup>d</sup> ± 0.10 |
| NB11 | 1.88<sup>e</sup> ± 0.07 | 1.87<sup>ab</sup> ± 0.34 |
| N20c | 5.92<sup>b</sup> ± 0.46 | 1.24<sup>bc</sup> ± 0.30 |
| N27d | 3.61<sup>d</sup> ± 0.21 | 0.73<sup>cd</sup> ± 0.00 |
| N29d | 1.94<sup>e</sup> ± 0.08 | 0.60<sup>cd</sup> ± 0.00 |

S.E. is standard error of the mean.
Means with same letter are not significantly different at $p \geq 0.05$ within columns using Tukey's test.

NB11 also presented comparably high N-fixing activity at 1.87 μg N mL^−1^ day^−1^ although producing fairly small halo on plate assay, yielding mean index value of 1.88. Strain N20c demonstrated second highest mean N-fixing index (5.92) but at fairly moderate amount of N-fixing activity (1.24 μg N mL^−1^ day^−1^). The lowest N-fixing activity was recorded in the range values of 0.34-1.38 μg N mL^−1^ day^−1^ by the rest of assessed isolates. Plate assays revealed lowest N-fixing index values in the range of 1.75-2.67. Although there was lack of reports on N-fixing activity (index) evaluation based on plate assay, the quantitative values were within range of other reported diazotrophic strains [40]. The ability of diazotrophic bacteria to convert atmospheric N_2_ into ammonia, the available form of N to be used by living organisms for biosynthesis is undeniably of vital importance to soil system especially through plants either through direct influence on plant growth, and indirectly through production of phytohormones that boost plant growth [50, 54].

Being one of the most limited but important plant growth nutrient, P mainly occurs in insoluble forms, whereby phosphate-solubilizing rhizobacteria plays a crucial role in converting it to available forms [55]. Among the 8 phosphate-solubilizing strains, PB3 and PB5 showed superior phosphate-solubilization index values in plate assay (4.21 and 4.10, respectively) whereas in enzymatic assay, the highest activity was demonstrated by isolate PC8 (307.7 μg mL^−1^) (Table 3).

**Table 3.** Phosphate-solubilization index and quantitative phosphate-solubilization estimation based on soluble P by phosphate-solubilizing isolates.

| Isolate | Phosphate-solubilizing index | Quantitative phosphate-solubilization (Soluble P in $\mu\text{g mL}^{-1}$) | Final mean pH in broth culture |
| --- | --- | --- | --- |
| | Mean value ± S.E. | | |
| PB5 | 4.21<sup>a</sup> ± 0.22 | 96.00<sup>d</sup> ± 2.65 | 4.27 |
| PC1 | 2.38<sup>cd</sup> ± 0.15 | 9.14<sup>f</sup> ± 0.75 | 4.93 |
| PB7 | 3.18<sup>b</sup> ± 0.23 | 159.00<sup>c</sup> ± 1.53 | 4.15 |
| PB3 | 4.10<sup>a</sup> ± 0.23 | 209.17<sup>b</sup> ± 8.97 | 4.11 |
| PC6 | 1.85<sup>ef</sup> ± 0.05 | 31.53<sup>e</sup> ± 0.39 | 4.63 |
| PC8 | 2.74<sup>bc</sup> ± 0.22 | 307.67<sup>a</sup> ± 8.41 | 4.04 |
| PB11 | 1.66<sup>f</sup> ± 0.08 | 3.25<sup>f</sup> ± 0.52 | 4.57 |
| PB6 | 2.19<sup>de</sup> ± 0.24 | 12.67<sup>f</sup> ± 0.55 | 4.20 |

S.E. is standard error of the mean.
Means with same letter are not significantly different at $p \geq 0.05$ within columns using Tukey's test.

The amount of phosphate-solubilization index and activities with final broth culture pH reported in this present study is comparable to the range reported by Pandey et al. [56], Gulati et al. [57], Song et al. [58]. Isolate PC8 showed higher phosphate-solubilization activity (307.67 μg mL^−1^) as compared with *Pseudomonas putida* (B0) at maximum activity of 247 μg mL^−1^ [56]. There was no significant relationship observed between phosphate-solubilizing index and soluble phosphate. Similar observations had been previously reported by Nautiyal [31] and Chang and Yang [59]. Nautiyal [31] stated the inability of phosphate-solubilizing *Pseudomonas aerogenes* and *Pseudomonas* sp.1 to produce halo on solid media but could solubilize substantial amount of P in different type of broth cultures. Nonetheless, there was a significant inverse correlation (r=−0.74, p ≤ 0.05) between soluble phosphate and pH of culture broth, and this observation corroborates previous findings [60–61]. The pH of broth declined to as low 4.04 as observed in PC8 activity from initial pH 7. This observation is consistent with earlier finding where there was a drop in pH due to bacterial activity is accompanied with mineral phosphate-solubilization and increment of its activity efficiency [56]. In their study on various organic acids secretion by 10 bacteria and 3 fungal strains, Rashid et al. [62] reported lowering of pH as part of the mechanism of microbial phosphate-solubilization.

### Cultural, biochemical, and molecular identification of selected functional bacterial isolates

The isolation of bacteria near the root zones of vegetations thriving in the rehabilitated forest generally lead to the recovery of putative beneficial rhizospheric or even free-living functional bacteria. In this present study, growth in the selective media proved to be a successful strategy to select bacteria based on their phenotypic activities on respective media. From the isolates assessed for their functional activities, 5 isolates of each functional group were selected to be further characterized on the basis of distinct colony morphology and high functional activities. After being selected based on these combined criteria, the isolates were grouped by morphological similarities and phenotypic characteristics.

On the basis of morphological characteristics of the 15 selected isolates depicted in Table 4, they exhibited varying properties. It was observed that the majority or 13 isolates exhibited round colonies in configuration or form; 10 colonies were convex whereas the rest shown raised elevations; 14 colonies were opaque in density; colonies of all of the isolates were almost creamy or yellowish in colour; 13 showed Gram negative reactions; all isolates were rods or short rods in single or in pairs arrangement; 13 without endospore production; and they were mostly motile as exhibited by 11 isolates. The following morphological properties presented more diverse characteristics: colony margin was represented by 8 entires, 3 undulates and serrates, respectively, and 1 lobate; colony size was represented by 6 moderates, 5 smalls, 3 larges and 1 pinpoint; and the shapes of endospores produced by 2 isolates were of rods and coccus, respectively.

**Table 4.** Morphological characteristics of selected functional isolates.

| Colony morphology | Isolate |  |  |  |  |  |  |  |  |  |  |  |  |  |  |
| --- | --- | --- | --- | --- | --- | --- | --- | --- | --- | --- | --- | --- | --- | --- | --- |
|  | CB15 | C22d | C29b | C41d | C46d | NB1 | NB10 | NB11 | N27d | NC9 | PB3 | PB5 | PB6 | PB7 | PC8 |
| **Configuration / Form** | Irregular | Round | Round | Irregular | Round | Round | Round | Round | Round | Round | Round | Round | Round | Round | Round |
| **Margin** | Undulate | Entire | Entire | Serrate | Entire | Entire | Undulate | Undulate | Entire | Entire | Entire | Serrate | Serrate | Entire | Lobate |
| **Elevation** | Raised | Convex | Convex | Raised | Convex | Convex | Raised | Convex | Convex | Convex | Convex | Raised | Raised | Convex | Convex |
| **Density** | Opaque | Opaque | Opaque | Opaque | Opaque | Opaque | Opaque | Opaque | Opaque | Opaque | Opaque | Opaque | Transparent | Opaque | Opaque |
| **Pigments** | Cream | Cream | Yellow | Cream | Cream | Cream | Cream | Cream | Yellowish cream | Cream | Yellowish cream | Cream | Cream | Cream | Cream |
| **Gram's reaction** | + | - | - | - | - | - | - | - | - | - | - | - | + | - | - |
| **Shape** | Rods | Rods | Rods | Short rods | Short rods | Short rods | Short rods | Rods | Short rods | Short rods | Rods | Rods | Short rods | Rods | Short rods |
| **Size** | Large | Moderate | Small | Large | Small | Pinpoint | Large | Moderate | Small | Moderate | Moderate | Moderate | Moderate | Small | Small |
| **Arrangement** | Single | Single | Single/pairs | Single | Single | Single | Single | Single | Single | Single/pairs | Single | Single/pairs | Single/pairs | Single/pairs | Single/pairs |
| **Endospore production** | + | - | - | - | - | - | - | - | + | - | - | - | - | - | - |
| **Endospore shape** | Rods | - | - | - | - | - | - | - | Coccus | - | - | - | - | - | - |
| **Motility** | + | + | + | - | + | - | + | - | - | + | + | + | + | + | + |

566 +, positive result; -, negative result.

The isolates were primarily identified based on molecular method, and in cases of multiple identities produced based on BLAST generated database, morphological (Table 4) and biochemical (Table 5) properties facilitated in the downsizing of identification task.

**Table 5.**
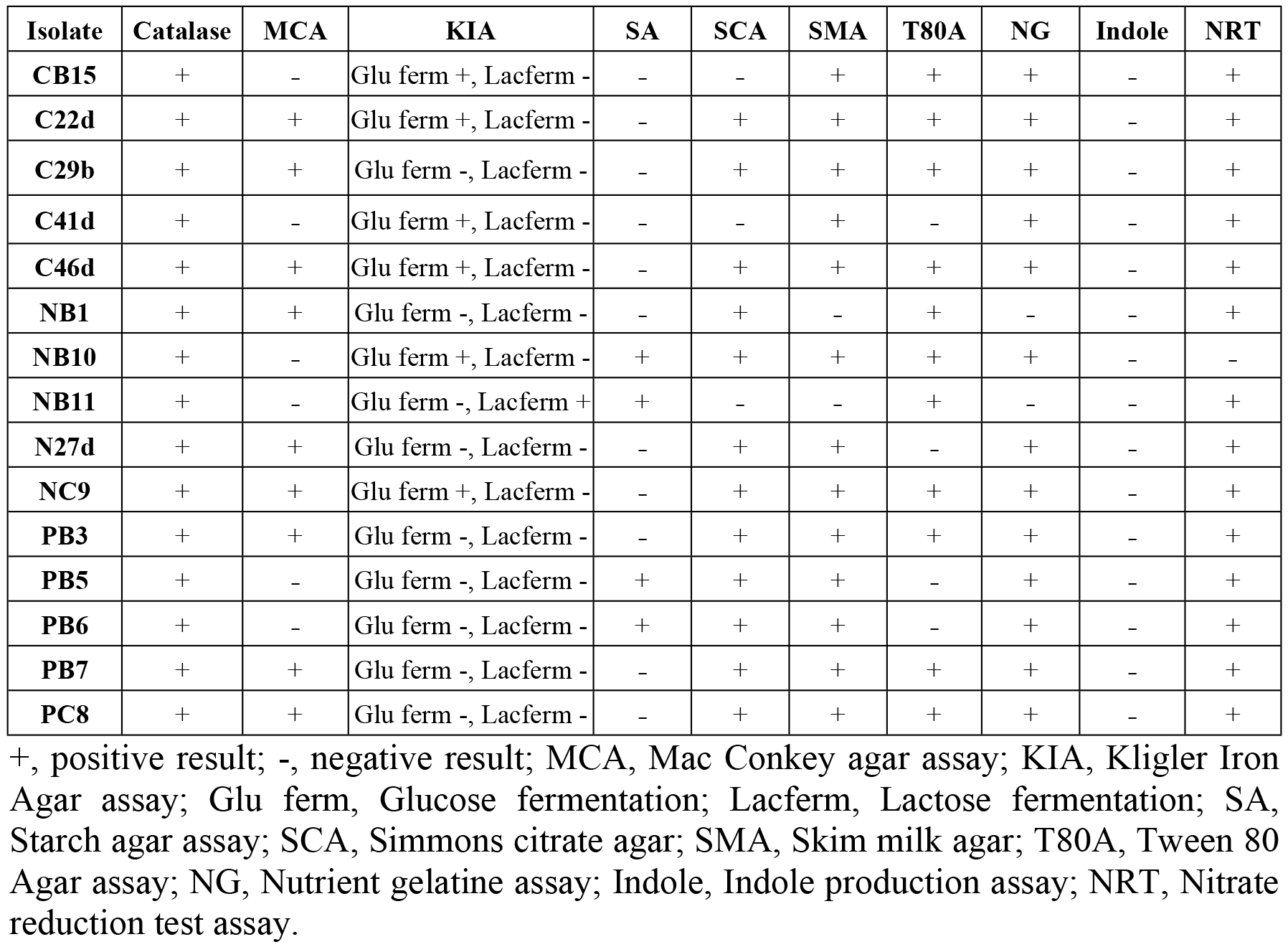
Biochemical properties of selected functional bacterial isolates.

In Table 6, the selected cellulolytic isolates in this present study were identified as *Serratia nematodiphila* strain SP6 (C46d), *Serratia marcescens* subspecies *sakuensis* isolate PSB23 (C22b), *Stenotrophomonas maltophilia* strain KNUC605 (C29b), *Bacillus thuringiensis* (Bt) (CB15), and *Stenotrophomonas* sp. Ellin162 (C41d). The N-fixing isolates were consisted of *Burkholderia nodosa* strain Br3470 (NB1), *Delftia lacustris* strain 332 (NC9), *Stenotrophomonas* sp. strain SeaH-As3w (N27d), *Ralstonia* sp. strain MCT1 (NB10), and *Cupriavidus basilensis* strain CCUG 49340T (NB11). The phosphate-solubilizing isolates were represented by *Burkholderia cepacia* strain 5.5B (PB5), *Burkholderia cepacia* strain ATCC 35254 (PB3), *Burkholderia uboniae* (PB7), *Burkholderia cepacia* strain 2EJ5 (PC8), and *Gluconacetobacter diazotrophicus* PAl 5 (PB6).

**Table 6.** Identities of cellulolytic, N-fixing, and phosphate-solubilizing bacterial isolates based on 16S rDNA analysis and BLAST databases.

| Phenotype | Isolate | Closest Identity | Similarity (%) | Accession No. |
| --- | --- | --- | --- | --- |
| Cellulolytic | C46d | *Serratia nematodiphila* strain SP6 | 98<br>(324/329) | JN315887.1 |
|  | C22b | *Serratia marcescens* subspecies *sakuensis* isolate PSB23 | 97<br>(578/594) | HQ242736.1 |
|  | C29b | *Stenotrophomonas maltophilia* strain KNUC605 | 99<br>(945/949) | HM047520.1 |
|  | CB15 | *Bacillus thuringiensis* | 99<br>(1018/1021) | FR846529.1 |
|  | C41d | *Stenotrophomonas* sp. Ellin162 | 100<br>(989/989) | AF409004.1 |
| N-fixing | NB1 | *Burkholderia nodosa* strain Br3470 | 98<br>(943/967) | AM284972.1 |
|  | NC9 | *Delftia lacustris* strain 332 | 99<br>(1009/1014) | EU888308.1 |
|  | N27d | *Stenotrophomonas* sp. SeaH-As3w | 100<br>(1029/1029) | FJ607350.1 |
|  | NB10 | *Ralstonia* sp. MCT1 | 99<br>(999/1007) | DQ232889.1 |
|  | NB11 | *Cupriavidus basilensis* strain CCUG 49340T | 99<br>(961/969) | FN597608.1 |
| Phosphate-solubilizing | PB5 | *Burkholderia cepacia* strain 5.5B | 99<br>(874/885) | FJ652617.1 |
|  | PB3 | *Burkholderia cepacia* strain ATCC 35254 | 99<br>(990/993) | AY741346.1 |
|  | PB7 | *Burkholderia uboniae* | 99<br>(993/997) | AB030584.1 |
|  | PC8 | *Burkholderia cepacia* strain 2EJ5 | 99<br>(1008/1016) | GQ383907.1 |
|  | PB6 | *Gluconacetobacter diazotrophicus* PAI 5 | 97<br>(778/801) | CP001189.1 |

These identities were matched with BLAST databases with the range of 97 to 100% base sequence similarities (Table 6). It was observed that identification of bacterial isolates over the three phenotypic groups revealed the predominance of Gram-negative over Gram-positive bacteria in the rehabilitated forest soils. The Gram-positive bacteria belong to only one taxonomic grouping, Firmicutes, represented by the genus *Bacillus*, whereas Gram-negative bacteria included Alpha-Proteobacteria with genus *Gluconacetobacter*; Beta-Proteobacteria represented by genera *Burkholderia, Delftia, Ralstonia*, and/or *Cupriavidus*; and Gamma-Proteobacteria by genera *Serratia* and *Stenotrophomonas*.

From the overall identification results of each functional group of isolates, bacteria from genus *Burkholderia* seemed to predominate in these tropical forest soils at one third of total identified isolates with distinguishing phosphate-solubilizing and N-fixing capabilities over other genera. *Stenotrophomonas* genus ranked second in abundance to *Burkholderia*. *Burkholderia* genus is a well-known classification of bacteria wherein its members possess vast versatility in niches and nutrition regulation such as N-fixing and phosphate-solubilization, as well as possessing numerous PGP attributes. *Burkholderia* species occur in the natural environment, even though some strains are also found in clinical samples, especially the *B. cepacia* complex species from patients with cystic fibrosis [63].

Based on our 16S rDNA sequences analysis results, the top 4 isolates with highest phosphate-solubilizing and N-fixing activities belong to genus *Burkholderia*, in which 3 of them were most closely related to *B. cepacia* (99% similarity) and *B. nodosa* (98% similarity), respectively. *Serratia marcescens* (of close similarity by C22b at 97%) or *Serratia* sp. (with C46d closely matched to *Serratia nematodiphila* by 98%) are known to have both cellulolytic and phosphate-solubilizing abilities [64–65]. Similar to the results of this present study, *Stenotrophomonas* spp. have been reported previously to possess both N-fixing [66] and cellulose-hydrolyzing abilities [67]. *Stenotrophomonas* species commonly prevail in nature, whereby *S*. *maltophilia* and *S. rhizophila* are reported to occur as rhizospheric and endophytic bacteria accountable for beneficial plant interactions [67]. The N-fixing isolates in this present study displayed higher diversity in terms of the species, as well as activities.

Diazotrophic isolate N27d revealed 100% matching identity with *Stenotrophomonas* sp. (strain SeaH-As3w) and it is believed to have cellolulytic potential similar to the C41d and C29b isolates. Previously, many studies have reported bacterial belonging to genus *Burkholderia* especially *B. cepacia* strains as having biocontrol ability against many phyto-pathogenic fungi such as *Fusarium* sp., *Pythium aphanidermatum*, *Pythium ultimum, Phytophthora capsici, Aspergillus flavus, Botrytis cinerea* and *Rhizoctonia solani* [68–72] as part of its main contributing PGP factors.

The phosphate-solubilizing bacteria showed the lowest diversity that only three species identities from two genera were recorded among the five identified isolates. Isolates PB3, PB5, PB7, and PC8 were closely matched (99% similarity) to *Burkholderia* sp. These isolates together with the N-fixing NB1 might potentially carry both the N-fixing and phosphate-solubilizing activities [73–74]. Diazotrophic isolate *Burkholderia nodosa* NB1 along with *Burkholderia mimosarum*, have been the latest classified novel species with N-fixing capacity and legume-nodulating bacteria that establish effective symbioses with legumes of *Mimosa* species [75–76]. All *Burkholderia* strains in this present study demonstrated variability in cultural and biochemical properties, which is typical for members of this genus.

The ability of *Burkholderia* strains to solubilize inorganic phosphates by various studies [77–80] have been well documented and nevertheless, a beneficial property in plant-growth promotion. Among the *Burkholderia* strains isolated in this study, this trait seemed most prevalent. Few *Burkholderia* strains are known for other PGP ability, which is to fix N_2_, either as free-living bacteria [81], or even nodulating ones that also fix N_2_ on media including *Burkholderia nodosa* strain as reported by da Silva et al. [82].

The genus *Delftia* (of close similarity by NC9 to *Delftia lacustris* at 99%) belongs to a newly grouped genus closely linked to *Comamonas*, in which *Delftia terephthalate* has been reported as a diazotrophic PGPR capable of degrading terephthalate, as well as antagonizing rice blast and blight causing *Xanthomonas oryzae, Rhizoctonia solani*, and *Pyricularia oryzae* [83–84]. One of the diazotrophic bacterium IHB B 4037 showing closely matched identity with *Delftia lacustris* isolated from rhizosphere of tea plant in India showed the highest nitrogenase activity in a study conducted by Gulati et al. [85]. *Delftia tsuruhatensis* was also revealed to be a diazotroph with biological control ability against several plant pathogens especially the three main rice pathogens in the likes of *Xanthomonas oryzae* pv. Oryzae, *Rhizoctonia solani*, and *Pyricularia oryzae* Cavara [86]. In summary, the variability among the bacterial strains of the same genera pointed to the fact that bacterial phenotypic properties are influenced by genetic, as well as environmental factors [87].

### Seed germination, shoot, and root elongation assays

Seed germination, shoot, and root elongation assays were done on 15 selected functional isolates on Green Gram seeds (Tables 7 - 9) to represent an attribute of early plant growth promotion activity by the isolates. After a week of growth, Green Gram roots originating from seeds treated with all selected bacterial strains exhibited different effects on root lengths, shoot lengths, and seedling vigor indexes compared with two controls, treated with distilled water and nutrient broth, respectively.

**Table 7.** Effects of selected cellulolytic bacterial inoculation on early growth stage of green gram.

| **Treatment** | **Mean number of seed germinated (3<sup>rd</sup> day)** | **Mean number of seed germinated (7th day)** | **Mean length of shoots (cm) (Mean ± S.E.)** | **Mean length of radicle (cm) (Mean ± S.E.)** | **% germinated seeds** | **Vigor index (Mean ± S.E.)** |
| --- | --- | --- | --- | --- | --- | --- |
| **C22b** | 50.0 | 50.0 | 12.84<sup>a</sup> ± 0.06 | 5.16<sup>ab</sup> ± 0.04 | 100 | 1799.50<sup>a</sup> ± 2.50 |
| **CB15** | 50.0 | 50.0 | 12.38<sup>a</sup> ± 0.45 | 5.47<sup>ab</sup> ± 0.75 | 100 | 1785.00<sup>ab</sup> ± 30.00 |
| **C41d** | 50.0 | 50.0 | 11.75<sup>a</sup> ± 0.03 | 5.67<sup>a</sup> ± 0.13 | 100 | 1741.50<sup>ab</sup> ± 10.50 |
| **C46d** | 50.0 | 50.0 | 11.73<sup>a</sup> ± 0.23 | 5.28<sup>ab</sup> ± 0.53 | 100 | 1700.00<sup>ab</sup> ± 30.00 |
| **C29b** | 50.0 | 50.0 | 11.81<sup>a</sup> ± 0.35 | 5.04<sup>ab</sup> ± 0.40 | 100 | 1684.50<sup>b</sup> ± 5.50 |
| **Control (Nutrient broth)** | 50.0 | 50.0 | 9.26<sup>b</sup> ± 0.18 | 3.36<sup>bc</sup> ± 0.02 | 100 | 1261.00<sup>c</sup> ± 19.00 |
| **Control (dH<sub>2</sub>O)** | 49.5 | 49.5 | 7.91<sup>b</sup> ± 0.18 | 2.87<sup>c</sup> ± 0.10 | 99 | 1067.00<sup>d</sup> ± 17.00 |

Data are presented as mean ± standard error (S.E.) of duplicates of 50 seedlings for each replicate per treatment for root elongation assay and seedling vigor index. The roots emerging from seeds failing to germinate after 2 days were previously marked and were not measured. The seedling vigor index (SVI) was calculated using formula: SVI = % of germination x seedling length (root length + shoot length). In the same column, significant differences according to Tukey's test at $p \leq 0.05$ level are indicated by different letters.

**Table 8.** Effects of selected N-fixing bacterial inoculation on early growth stage of green gram.

| Treatment | Mean number of seed germinated (3<sup>rd</sup> day) | Mean number of seed germinated (7<sup>th</sup> day) | Mean length of shoots (cm) (Mean ± S.E.) | Mean length of radicle (cm) (Mean ± S.E.) | % germinated seeds | Vigor index (Mean ± S.E.) |
| --- | --- | --- | --- | --- | --- | --- |
| **NB1** | 50.0 | 50.0 | 11.27<sup>a</sup> ± 0.06 | 3.84<sup>ab</sup> ± 0.02 | 100 | 1510.00<sup>a</sup> ± 4.00 |
| **NC9** | 50.0 | 50.0 | 11.00<sup>a</sup> ± 0.32 | 4.10<sup>a</sup> ± 0.34 | 100 | 1510.00<sup>a</sup> ± 66.00 |
| **NB10** | 50.0 | 50.0 | 10.81<sup>ab</sup> ± 0.15 | 3.89<sup>ab</sup> ± 0.24 | 100 | 1469.00<sup>a</sup> ± 9.00 |
| **NB11** | 50.0 | 50.0 | 10.43<sup>ab</sup> ± 0.27 | 4.09<sup>a</sup> ± 0.26 | 100 | 1451.50<sup>ab</sup> ± 52.50 |
| **N27d** | 50.0 | 50.0 | 10.31<sup>ab</sup> ± 0.61 | 3.68<sup>ab</sup> ± 0.19 | 100 | 1399.00<sup>ab</sup> ± 42.00 |
| **Control (Nutrient broth)** | 50.0 | 50.0 | 9.26<sup>bc</sup> ± 0.18 | 3.36<sup>ab</sup> ± 0.02 | 100 | 1261.00<sup>bc</sup> ± 19.00 |
| **Control (dH<sub>2</sub>O)** | 49.5 | 49.5 | 7.91<sup>c</sup> ± 0.18 | 2.87<sup>b</sup> ± 0.10 | 99 | 1067.00<sup>c</sup> ± 17.00 |

Data are presented as mean ± standard error (S.E.) of duplicates of 50 seedlings for each replicate per treatment for root elongation assay and seedling vigor index. The roots emerging from seeds failing to germinate after 2 days were previously marked and were not measured. The seedling vigor index (SVI) was calculated using formula: SVI = % of germination x seedling length (root length + shoot length). In the same column, significant differences according to Tukey's Test at $p \leq 0.05$ level are indicated by different letters.

**Table 9.**
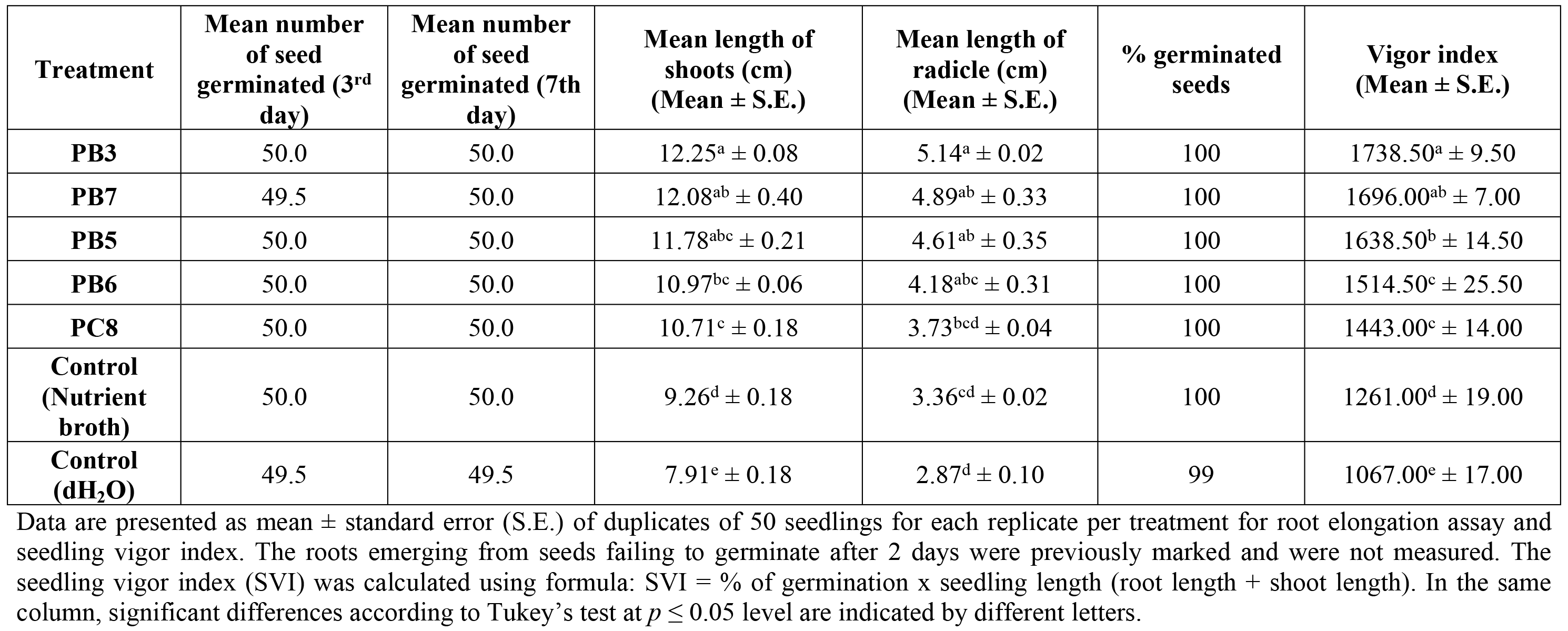
Effects of selected phosphate-solubilizing bacterial inoculation on early growth stage of green gram.

As shown in Table 7, inoculation with cellulolytic isolate C41d (*Stenotrophomonas* sp.) resulted in significantly longer root lengths compared to both control treatments, but was not significantly different from the rest of cellulolytic isolates. On the other hand, the treatments with all selected cellulolytic strains resulted in significantly higher shoot lengths compared to both controls but not different with one another, which ranged from 11.46 to 12.9 cm. In terms of SVI, all cellulolytic strains exhibited significantly higher values than both controls, with isolate C22b (*Serratia marcescens* subsp. *sakuensis*) presented outstanding values but was not different with isolates CB15, C41d, and C46d (Bt, *Stenotrophomonas* sp., and *Serratia nematodiphila*) which ranged from 1670 to 1815.

In Table 8, the treatments with N-fixing isolates NC9 (*Delftia lacustris*) and NB11 (*Cupriavidus basilensis*) showed higher root lengths and shoot lengths occurring in the range of 3.76 cm to 4.44 cm and 10.68 cm to 11.33 cm, respectively, whereas SVI values for isolates NB1, NC9, and NB10 (*Burkholderia nodosa, Delftia lacustris*, and *Ralstonia* sp.) revealed significantly higher SVI values than both controls, ranging from 1444 to 1576, but were not significantly different from the rest of N-fixing strains.

Isolates PB3, PB5, and PB7 (*Burkholderia cepacia, Burkholderia cepacia*, and *Burkholderia uboniae*) resulted in the longest root lengths among the phosphate-solubilizing strains in the range of 4.26 cm to 5.16 cm and SVI in the range of 1624 to 1748 using these three isolates (Table 9). Significantly higher shoot lengths however, were achieved with phosphate-solubilizing isolates than both controls, ranging from 10.53 cm to 12.33 cm. Isolate PB3 had the most outstanding effect on shoot length, but not different from those of isolates PB5 and PB7, among other phosphate-solubilizing bacterial strains.

A previous study by Satya et al. [88] confirmed similar observation on *Burkholderia* sp. as their application using *Burkholderia* sp. strain TNAU-1 demonstrated improvement in root and shoot lengths, as well as increased seedling vigour on mung bean. According to Egamberdieva et al. [89], inoculation of one of the salt tolerant rhizobacterial strains, *Stenotrophomonas rhizophila* ep-17 resulted in improved dry weight of cucumber by up to 68% in various salt concentrations as high as 10 dSm^−1^ in comparison to non-inoculated plants exposed to salt stress.

Hameeda et al. [90] demonstrated that one of phosphate-solubilizing bacterial strains in the study, *Serratia marcescens* designated as EB 67 tested on maize growth by paper towel method was able to produce higher root and plumule lengths, plant weight (43%), and seed vigor index, whereas another strain also included in the study, *Serratia* sp. designated as EB75 resulted in increased root length, plant weight by 23%, and seed vigor index over uninoculated control.

Kang et al. [91] reported that inoculation of a novel cold stress resistant rhizobacteria *Serratia nematodiphila* PEJ1011 on pepper plants resulted in the latter’s significantly improved growth by means of both increased root and shoot lengths, and fresh weight of the peppers on both normal and cold temperature as compared with controls under normal and cold temperature, respectively. Most previous studies on Bt reported it as a safe and effective biological control agent for the suppression of a broad type of insect pests [92–95] due to its ability to produce parasporal crystals consisting of protein molecules called delta endotoxins which are toxic to insect pests [96]. Besides its apparent biological control attribute, current study by Khan et al. [97] also reported plant growth promotion *via* increase in plant growth parameters on mung bean seeds coated with various Bt strains used in the study. In their study, all the tested strains improved shoot weight, with strain BT-64 in particular, was reported to have resulted in 97%, 42%, 46%, and 71% increment in shoot weight, shoot length, root length, and root weight, respectively over control.

Apart from having high nitrogenase activity strain from rhizospheric soil of tea plants [85], Jin et al. [98] found that *Delftia lacustris* isolated from rhizospheric soils of tobacco to be one of strongly antagonistic bacteria against pathogen *Phytophthora nicotianae* which could have potentially helped promoting growth of tobacco crop. According to a study by Yrjȁlȁ et al. [99], *Cupriavidus basilensis* of Burkholderiaceae family, and especially strains of the genus *Cupriavidus* have been shown to be isolated from hybrid aspen root in mostly pristine soil as compared to rhizospheric soils contaminated with polyaromatic hydrocarbon (PAH) constituents.

On the other hand, the addition of PAH mixture (anthracene, phenanthrene, and pyrene) resulted in the apparent shift in the diversity of uncultured *Burkholderia* in rhizosphere of hybrid aspen and lowest fraction (five out of 38 sequences) of this genus was found in pristine rhizosphere. Their study indicated the main difference in terms of soil diversity that could influence the abundance between both genera. *Cupriavidus basilensis* designated HMF14 has also been recently identified as an important furan-degrading bacterium, and was isolated by employing an elegant assay based on the germination of lettuce seeds [100]. Furanic compounds as toxic fermentation inhibitors are derived from lignocellulose hydrolysates in which occurrences of the former in high amount could pose carcinogenicity effects to humans [100].

*Cupriavidus* (synonym *Ralstonia*) is closely linked to *Burkholderia*, but compared to the latter genus it consists of very limited N-fixing species [101]. Both genera, in particular consisting of *Burkholderia* spp. and *Cupriavidus taiwanensis* have been reported to be N-fixing symbionts of legume *Mimosa* with the ability to nodulate the latter [102–104]. Some strains of *Burkholderia* have been known as Beta-rhizobia which could exclusively nodulate large legume of genus *Mimosa* (Mimosoideae) [101]. These legume-nodulating burkholderias are not closely related to phytopathogenic or pathogenic group of burkholderias including the *Burkholderia cepacia* complex, but instead are closely linked to PGPR, mostly diazotrophic, endophytic environmental *Burkholderia* species and strains [105].

*Burkholderia nodosa* designated as LMG 23741 has also been isolated from acidic soils of Amazon which was a nodulating N-fixing bacterium [82], as compared to the strain in this present study which did not nodulate, but was a free-living or rhizospheric N fixer. Both strains are known to be absent in ability to solubilize inorganic phosphates. The strain reported by da Silva et al. [82] also did not possess anti-fungal characteristic. Besides the N-fixing isolate *Ralstonia* sp. characterized in this present study, *Ralstonia taiwanensis* is the first N-fixing beta-proteobacterium strain having the ability to nodulate the roots of *Mimosa* from which it was isolated [106]. However, some *Ralstonia* spp. are environmental variant microorganisms and there are also phytopathogenic strains such as *Ralstonia solanacearum* that cause bacterial wilt occurrence on a broad range of crops [107].

*Burkholderia cepacia* designated as LMG 1222 by da Silva et al. [82] was also reported as a non N-fixing bacteria isolated from acidic Amazon soils which could solubilize inorganic phosphate, as well as possessing anti-fungal property. However, there is lack of literature on beneficial properties of *Burkholderia uboniae* and hence, the results of this present study could contribute primary evaluation on the potential of this particular strain on early plant growth promotion. Many studies reported bacterial belonging to genus *Burkholderia* especially *B. cepacia* strains as having biocontrol ability against many phyto-pathogenic fungi such as *Fusarium* sp., *Pythium aphanidermatum*, *Pythium ultimum, Phytophthora capsici, Aspergillus flavus, Botrytis cinerea*, and *Rhizoctonia solani* [68–72] as part of its main contributing PGP factors. *Gluconacetobacter diazotrophicus* is one of diazotrophic species reported to form anomalous symbiotic relationship with plants by thriving on the root system surface as rhizobacteria, whereas a number of strains are known to infect plant tissues as endophytic bacteria and fix N_2_, resulting in enhanced plant growth [108–111].

### Indole-3-acetic acid assay

On the basis of respective phenotypic functional activities, Green Gram seed germination, and root elongation assay results, 7 bacterial isolates were selected for IAA production assay (Table 10).

**Table 10.** Indole-3-acetic acid production by selected functional bacterial isolates.

| Phenotype | Isolate | IAA production<br>($\mu\text{g mL}^{-1}$)<br>Mean ± S.E. |
| --- | --- | --- |
| Cellulolytic | *Stenotrophomonas maltophilia* C29b | 12.38<sup>a</sup> ± 1.48 |
| | *Serratia nematodiphila* C46d | 11.39<sup>ab</sup> ± 1.68 |
| | *Bacillus thuringiensis* CB15 | 0.98<sup>c</sup> ± 0.16 |
| Phosphate-solubilizing | *Burkholderia cepacia* PC8 | 8.45<sup>ab</sup> ± 0.28 |
| | *Burkholderia cepacia* PB3 | 7.81<sup>b</sup> ± 0.11 |
| | *Gluconacetobacter diazotrophicus* PB6 | 7.51<sup>b</sup> ± 0.17 |
| Nitrogen-fixing | *Burkholderia nodosa* NB1 | 0.43<sup>c</sup> ± 0.45 |

S.E. is the standard error of the mean value.
Significant differences according to Tukey's test at $p \leq 0.05$ level are indicated by different letters.

All the 7 selected strains produced IAA without supplementation of L-tryptophan, from N-fixing isolate *Burkholderia nodosa* NB1 to most outstanding IAA production by cellulolytic isolates *Stenotrophomonas maltophilia* C29b and *Serratia nematodiphila* C46d, and phosphate-solubilizing isolate *Burkholderia cepacia* PC8 ranging from 8.17 to 13.86 μg mL^−1^. The IAA produced by the cellulolytic, non N-fixing *Stenotrophomonas maltophilia* strain in this present study, C29b was at 12.38 μg mL^−1^ with absence of tryptophan. *Stenotrophomonas maltophilia* has been previously reported as diazotrophic with IAA production of 2.6 mg mL^−1^ [112], 112.8 μg mL^−1^ for S. *maltophilia* PM-1, and 139.2 μg mL^−1^ in S. *maltophilia* PM-26 [113] with L-tryptophan as supplementation.

*Serratia nematodiphila* C46d strain in this present study was found to synthesize IAA at 11.39 μg mL^−1^. Previously, *Serratia nematodiphila* designated NII-0928 isolated from Western ghat forest soil was found to possess multiple PGP attributes including phosphate-solubilization and IAA production at higher concentration, at 76.6 μg mL^−1^ and 58.9 μg mL^−1^, respectively [114]. As stated previously, the variability among the bacterial strains of the same genera points to the fact that bacterial phenotypic properties are influenced by genetic and environmental factors [87]. *Serratia nematodiphila* LRE07 has also been reported to be one of the heavy metals-resistant endophytic bacteria isolated from *Solanum nigrum* L. planted in metal-polluted soil which improved plant growth and Cd extraction from soil as phytoremediation activity [115]. *Bacillus thuringiensis* (Bt) strain CB15 in this present study yielded IAA amount within the stated range as reported by Vidal-Quist et al. [116], whereby all of 44 Bt strains screened in their studies were reported to produce IAA in a range of 0.12 to 6.59 μg mL^−1^. The IAA production range is comparable to the findings of Raddadi et al. [117].

Besides being well known for having biocontrol activity against insects [118], phytopathogens including fungi [119–120], and bacteria [121], Bt has the potential to promote plant growth *via* production of phytohormones, such as auxins, or by regulating levels of plant ethylene [117, 122], stimulation of nodulation in legumes by some strains and hence, *Rhizobium-based* N_2_ fixation [123–124], improvement by other strains of mineral nutrients (such as Fe and P), recovery by the plant [117], and promote protection of plants from abiotic stresses [125].

*Burkholderia cepacia* strains PC8 and PB3 evaluated in this present study without addition of tryptophan produced IAA at 8.45 and 7.81 μg mL^−1^, respectively. Bevivino et al. [126] reported that rhizospheric strains of *Burkholderia cepacia* PHP7 and TVV75 were found to produce IAA at approximately 0.6 and 0.4 μg mL^−1^, respectively, both of which were far lacking by at least 14-fold, even with addition of L-tryptophan, in comparison to strains in this present study. This phenomenon could indicate that the *B. cepacia* strains were actively involved in IAA synthesis in pure culture [127].

Phosphate-solubilizer and non N-fixing *Gluconacetobacter diazotrophicus* strain PB6 produced IAA at 7.51 μg mL^−1^ without precursor tryptophan, which was higher than those reported previously and improved shoot length and vigor index of green gram than uninoculated controls. Anitha and Thangaraju [128] reported an endophytic diazotroph *Gluconacetobacter diazotrophicus* strain in their study to have exhibited nitrogenase activity, phosphate-solubilization, and IAA production with and without tryptophan at 4.74 μg mL^−1^ and 1.4 μg mL^−1^, respectively. The strain in their study could also colonize roots of paddy in the growth solution under aseptic hydroponic condition. The researchers further stated that inoculation of rice seedlings with the strain also resulted in improved shoot length, root length, and plant biomass than uninoculated control.

Despite the least amount of IAA produced by *Burkholderia nodosa* strain NB1 at 0.43 μg mL^−1^, this is the first results on IAA production ever reported for non-nodulating N-fixing *Burkholderia nodosa* strain. In fact, there is dearth of reports concerning the beneficial PGP traits of *Burkholderia nodosa* with exception of it being reported as N-fixing symbiont of nodulating legumes by being isolated from N-fixing root nodules of legumes *Mimosa bimucronata* and *Mimosa scabrella* [129], and its apparent ability as biocontrol agent against bacterial wilt of tomato, as well as *Fusarium* wilt of tomato and spinach with recorded suppression of disease index by 33-79% [130].

The presence of an appropriate precursor such as L-tryptophan may mediate the production of auxins by some microorganisms [131]. The influence of auxins on plant seedlings relies upon their concentrations. For instance, high concentration may suppress growth, whereas low concentration may produce stimulative effects on growth [132]. The absence of tryptophan precursor in IAA production has also been previously observed in other bacterium, *Azotobacter brasilense* in a mechanism known as a tryptophan-independent pathway which was suggested during experiments using labelled tryptophan [133].

Despite this fact, there is still absence of clear genetic or biochemical proof to be established to consolidate this pathway [134]. According to Spaepen and Vanderleyden [134], occurrence of some IAA production despite inactivation of a single pathway on some knockout studies revealed redundancy of IAA biosynthetic pathways in some studied microorganisms, indicating that several pathways do exist and operating in a single microorganism. Nevertheless, the ability to secrete IAA without absolute dependence on tryptophan as the main precursor proved to be an added advantage over tryptophan dependent microbes in plant rhizospheres to produce beneficial associative effects with plant roots to enhance plant growth.

### Conclusions

Out of the 15 identified, bacterial isolates possess beneficial phenotypic traits, a third belong to genus *Burkholderia* and a fifth to *Stenotrophomonas* sp. with both genera consisting of members from two different functional groups. Among the tested bacterial strains, isolate C46d, NB1, and PC8 showed outstanding cellulase, N-fixing, and phosphate-solubilizing activities, respectively. The results of the experiments confirmed the multiple PGP traits of the selected bacterial isolates based on their respective high functional activities, root, shoot lengths, and seedling vigour improvements when bacterized on mung bean seeds, as well as presented some significant IAA productions. All these functional bacterial strains could potentially produce beneficial synergistic effects *via* their versatile properties on improving soil fertility and possible plant growth stimulation in beneficial interactions. Further characterizations of these potent strains will be helpful in elucidating their potential uses in biotechnological applications.

## Acknowledgements

Many thanks are extended to the colleagues and staff of Universiti Putra Malaysia Bintulu Sarawak Campus and Department of Agriculture, Kuching, Sarawak for their valuable cooperation and support. Contribution of the late Dr. Wong Sing King was also greatly appreciated for his advice on microbiological aspects of the study.

